# GePI: Retrieval of fully automated recognition and extraction of gene and protein interaction mentions from unstructured literature

**DOI:** 10.1101/2022.07.08.499305

**Authors:** Erik Faessler, Udo Hahn, Sascha Schäuble

## Abstract

**Motivation:** Knowledge about interactions between genes and proteins is vital for bio-molecular research. A large part of this knowledge is published in written text and not accessible in a structured way. To remedy this situation, several repositories of automatically extracted interaction facts were proposed over the years. However, existing solutions lack key features such as permanently updated data resources, easy accessibility and structured result generation ready to be used for downstream analyses.

**Results:** We propose GePI, a database portal for fully automated extraction and presentation of molecular interaction facts from scientific literature. GePI offers batch queries, immediate inspection of textual evidence and full text filters. To this end, GePI leverages two gene recognition and normalization approaches as well as optimized runtime for molecular event extraction. The resulting natural language processing pipeline is applied to the full set of publicly available documents from PubMed and the PubMed Central open access subset accounting for more than 33M abstracts and 4.2M complete articles as of 2022. To accommodate the rapid growth of the scientific literature, the fact database is automatically updated several times per week. In summary, our web application GePI allows for the first time a free and easy-to-use investigation of gene and protein interaction information as soon as they are published with unique query possibilities.

**Availability and Implementation:** The GePI web interface is available at http://gepi.coling.uni-jena.de.

## 1 Introduction

The molecular interactions between genes or gene products are the subject of a large body of scientific research that is constantly growing. Without structured databases it is already challenging to gain a comprehensive overview of already described functional or physical associations between a particular set of genes or proteins. Towards the goal of creating such resources, several curated databases on protein-protein interactions have been developed. Example databases include IntAct (Orchard *et al*., 2014), BioGrid (Oughtred *et al*., 2021) and HPRD (Keshava Prasad *et al*., 2009) that each contain millions of interactions curated from thousands of publications. See Lehne and Schlitt (2009) for a thorough overview. Such databases offer detailed, cleanly assembled interaction data and by doing so enable continuing scientific hypothesis testing and discovery. However, only a fraction of the complete set of PubMed with over 33M citations and PubMed Central (PMC) with more than 4.2M full texts in its open access subset as of 2022 is covered by such structured databases. Given the ever accelerating speed of publication generation, the gap in available knowledge and its structured representation in domain-specific databases will likely only increase. Therefore, automatic means of scanning the biomedical literature have gained attention over the years. String (Szklarczyk *et al*., 2021) assembles evidence for protein-protein interactions from existing databases and computational interaction predictions and also contains an estimation of protein-protein associations by weighted co-occurrence measures based on an automated literature scan. Fueled by the efforts of the BioNLP Shared Task (ST) 2009 and following challenges (Kim *et al*., 2009, 2011; Nédellec *et al*., 2013), software tools for the extraction of high-quality molecular events became publicly available. In part these were integrated into databases like BioContext (Gerner *et al*., 2012) and EvexDB (Van Landeghem *et al*., 2013) that allow exploring automatically extracted molecular associations for one gene or protein at a time in the web application or via API access. These tools have the common goal to offer aggregated knowledge from the literature in a user-friendly way that would otherwise require an extensive, perhaps even prohibitively large amount of manual effort.

Despite the progress towards this goal, existing tools come with a number of limitations. Most relevant web sites support the exploration of the molecular interaction literature by specifying single genes or proteins for subsequent manual inspection. Given the rise of next-generation sequencing techniques that rapidly provide a multitude of data, e. g. sets of differentially expressed genes, large lists of gene or protein identifiers are common user inputs. Consequently, the demand for the retrieval of molecular associations from the literature on basis of extensive input lists needs to be addressed. Additionally, filter capabilities on the document context of retrieved interactions or on the type of full-text sections are unavailable in existing databases. However, expressive filters are essential to provide interaction descriptions that occur in a specific context such as the mention of a disease. Filters on section headings allow to restrict the interaction retrieval to, e. g., results sections, but not the abstract, of full texts. Also, existing systems became unavailable or outdated over time. To fill this gap, we here present Gene and Protein Interactions (GePI), a tool for the fully automated and constantly updated batch retrieval of interaction mentions in the scientific literature. GePI can be used to find all interaction mentions for a predefined gene/protein list to other genes or to find all interaction mentions between members of two provided lists. Next to filtering for specific interaction types, the web application offers full text search on the sentence and paragraph levels surrounding an interaction mention as well section filtering and a download of all found interactions with aggregated statistics as an Excel spreadsheet file.

## 2 System and methods

Figure 1 shows the components of the GePI application ecosystem. Gene information is modelled in a Neo4j graph database (see subsection 2.3 for details). These information are incorporated in the natural language processing (NLP) pipeline for the resolution of gene groups, families and complexes. The input to the pipeline are XML documents from PubMed and PMC. In the NLP pipeline mentions of genes, gene products and molecular events between those are extracted and sent to an ElasticSearch (ES) cluster for indexing. Finally, the web application leverages the ES index and the gene database to serve use requests.

**Fig. 1.**
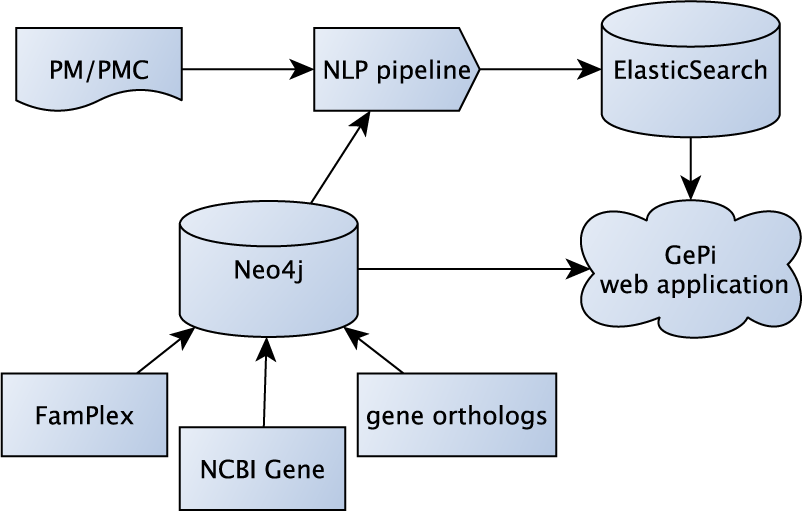
Illustration of the GePI components and their inner dependencies.

### 2.1 NLP Pipeline

To shape the ES index to GePI’s requirements, a series of processing steps are conducted for the input PubMed (PM) and PMC documents. These steps include the recognition and extraction of the mentions of molecular events between genes or proteins in any given sentence of a scientific text. We additionally assembled and assigned unique database identifiers from NCBI Gene to gene and protein mentions. These served to create canonical links between the entries of the NCBI Gene database and the literature and allowed to efficiently resolve lexical diversity and ambiguity of gene and protein names.

As depicted in Figure 2, the NLP pipeline encompasses components for basic segmentation and syntactic analysis tasks as well as semantic analysis for genes and gene products. We used two different algorithmic approaches for gene mention recognition and their normalization into NCBI Gene, namely GNormPlus and GeNo. This may lead to conflicting gene annotations where the begin and end of multiple gene mentions overlap but are neither equal nor disjunct. In case of overlapping gene/protein mentions, the result of both tools are merged on two levels: Firstly, overlapping gene or protein mentions are collapsed into the longest of the mentions. The reasoning behind this heuristic is that longer mentions may contain more specifiers (e. g. *AMPK* vs. *AMPK α 1*) and is thus more concrete. Secondly, all identified IDs belonging to a gene or protein from an overlapping set of mentions are assigned to the unified mention identified in step 1. Gene or protein mentions without overlap are directly put into our index database. Entities tagged as gene or protein families, groups or protein complexes are kept for grounding into a database of gene and protein families and complexes (see subsection 2.2). Such mentions are frequently omitted by gene normalization algorithms since they cannot be grounded into databases about individual genes such as NCBI Gene or UniProt. Still, such entities may participate in the description of molecular interaction events in the literature.

**Fig. 2.**
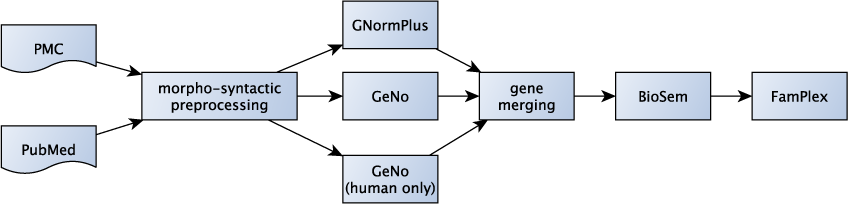
Schematic display of the gene and gene product event extraction pipeline used for GePI.

Event extraction is performed by BioSem, an efficient approach based on the automatic learning of rules from example data (Bui and Sloot, 2012; Bui *et al*., 2013). All gene and protein mentions that result from the merging operation are potential participants in interaction events found by BioSem, i. e. it is possible to find interactions between entities recognized by different systems. BioSem analyses document text sentence-wise. Consequently, all molecular interactions are contained by a single sentence in GePI. In their original form, the output events are tree structures where events can also be arguments of other events. To represent strictly binary interactions that facilitate querying, display and statistical description of the interaction data, complex events are broken down to combinations of argument pairs that consist solely of gene or protein mentions. Consequently, GePI’s interaction database consists of index items that have exactly two gene or protein interaction partners.

In the last step, the extracted events are sent to ES for indexing. Information about the mention and ID source – i.e. which NER and normalization component produced the respective datum – is stored with the index items for display and filter facilities in GePI. This pipeline is applied automatically to PubMed (PM) and PubMed Central (PMC) updates several times per week to offer up-to-date access to the molecular interaction literature. Its components are available in the JCoRe (Hahn *et al*., 2016) component repository with the exception of GNormPlus.

### 2.2 Gene Normalization

For the unambiguous identification of genes and gene products in the processed literature we employ gene normalization algorithms. The task of such software is to assign a unique identifier from a database (termed *mapping* or *grounding*), in this case NCBI Gene, to each gene mention in the document texts. The normalization step is critical for the core functionality of GePI: the query of interactions by lists of gene or protein identifiers. The effects of the normalization are two fold. On the query side, the lexical variability of gene and protein names is accounted for. The gene ID serves as a single point of reference independently of the actual lexical form a gene or gene product was referred to in the text. It is therefore not necessary to specify synonyms as input of GePI to capture all occurrences of a gene or protein in the literature. On the literature side, gene normalization resolves ambiguity that arises from homonymy. Some gene or protein names denote multiple, separate items in gene and protein databases. For example, the name *ARP5* is a synonym for the NCBI Gene symbols *ACTR3, ANGPTL6* and *APOBEC3C*. In PubMed the *actin-related protein* meaning of the name dominates the literature but is also used to denote *angiopoietin-related growth factor protein 5* in some cases. Dependent on the document context, the ID of the gene that was most probably referred to by the author must be assigned to avoid the dilution of GePI results by undesired gene items sharing the same synonym.

In GePI, a total of three gene normalization algorithms were employed. We chose GNormPlus for its high scores in human and intra-species gene normalization tasks (Wei *et al*., 2015). However, GNormPlus uses conditional random field (CRF)s based on manually extracted text features in its entity recognition module. gene recognition (GR) approaches based on deep learning (DL) show superior annotation quality in a large range of biomedical annotation tasks even for non-specialized DL models (Habibi *et al*., 2017). Additionally, the resources used by GNormPlus are several years old. This is problematic due to the fact that no facilities for the update of the gene database are available. An up-to-date gene database is required for access of newly described genes or proteins in the literature.

For the aforementioned reasons, we decided to update our GeNo system (Wermter *et al*., 2009) to alleviate these shortcomings and thus complement GNormPlus. The original GeNo system revealed a high evaluation score on the BioCreative II (BC2) gene normalization (GN) challenge. Since this challenge was restricted to the normalization of human gene mentions, GeNo was not directly applicable to the multi-species gene normalization task in a general retrieval application like GePI. Similar to GNormPlus, GeNo used a feature-based CRF algorithm for GR as was the state-of-the art at the time. Thus, the update to GeNo encompasses the incorporation of an up-to-date DL-based GR algorithm (described in Faessler *et al*. (2020)), a gene name composite resolution module, a species-assignment module and a fully automatic gene database creation algorithm with virtually no restrictions on the included species. To accommodate for medical research, we additionally added a version that is restricted to mentions of human genes. In this case, all gene mentions are assigned human gene IDs.

In their original form, both GN tools recognized mentions of individual genes and their products as well as groups or families of them. Due to their adaption to GN tasks, they suppress the output of family names. Regarding molecular interaction extraction, however, families and protein complexes might be arguments of textual statements in the literature. Because of their importance in the description of biomolecular interactions, we decided to keep discarded gene and protein group, family and complex mentions. This required a few alterations to the source of code of GNormPlus.

The incorporation of new biomolecular objects leads to new challenges regarding entity grounding. Common gene and protein databases like NCBI Gene or UniProt list the individual building blocks of protein complexes but do not contain entries for the resulting objects itself. This may lead to a new source of normalization errors. For example, the 5’ adenosine monophosphate-activated protein kinase (AMPK) enzyme consists of three subunits, *α, β* and *γ*. Each of those subunits is part of a family of two to three isoforms. For each of these isoforms, entries in NCBI Gene and UniProt exist. A common mapping mistake is to ground mentions of the AMPK complex to one of its isoforms. While this is an adequate assessment - the subunits are implicitly present when their complex is mentioned - a structurally more comprehensive method were to include a) a normalized denotation of the complex itself and b) the additional tagging of all three subunits on the same lexical unit. These requirements have been formerly identified and led to the development of the FamPlex resource by Bachman *et al*. (2018). It lists gene and protein families and complexes with diverse lexical synonyms for the textual grounding of the resource entries into the literature. Additionally, isoforms, families and complexes are connected via *IS-A* and *PART-OF* relations that represent the tree structure induced by the different description levels of protein complexes. We incorporated this resource by matching the entity strings specified in FamPlex to entities tagged as families or protein complexes by GNormPlus or GeNo via string matching. We mapped the leafs of the tree structures - the individual genes or proteins - to NCBI Gene and therefore anchored them in the text documents via the employed GN algorithms. This enables users to issue GePI queries for protein complexes and families that otherwise would yield empty results.

### 2.3 Gene Data Model

The gene data basis for GePI is an integration of NCBI Gene, the NCBI gene ortholog file (Agarwala *et al*., 2018) and FamPlex. We use NCBI Gene as a flat list of data records that is a source of symbols, names, synonyms and descriptions for each entry. *Gene orthologs* assembles individual NCBI Gene records to an abstract concept of functionally similar genes across different species. Each ortholog group is materialized in our data model with connections to its NCBI Gene elements. In a similar fashion, FamPlex organizes sets of gene records to families and complexes, sometimes across multiple levels of *IS-A* and *PART-OF* relations. Where applicable, FamPlex specifies links to external resources that we used to foster an attachment to NCBI Gene to obtain a fully integrated gene data model including orthologs, families and complexes.

### 2.4 Interaction Database

The GePI interaction database contains one record for each gene/protein interaction that was extracted from the literature. The records contain exactly two gene or protein arguments which is the central interaction information. The entities are tagged with multiple IDs for different ways to obtain them later. We store the mention text itself for fallback text-based queries, the NCBI Gene ID, its internal data model ID and the data model IDs of all directly or indirectly governing data model concepts such as orthologs, families and protein complexes. In this way, each interaction record can be retrieved by queries aiming at the different data abstraction levels. The interaction partners can be found directly by searching for their NCBI Gene ID but also by queries containing the ID ortholog group the interaction partners belong to, for example.

The records include the sentence and paragraph texts in which the interaction was found as well as the headings of all sections and subsections the interaction lies in. These information enable the full-text search and filter features of GePI. Since BioSem works on the sentence level, there is exactly one sentence per interaction record. The paragraph is one of three text scopes. On the abstract level, it is either the whole abstract or, when the abstract is structured, the respective section of the abstract like *Background, Methods, Results* etc. In PubMed Central (PMC) full texts, the paragraphs are encoded directly in the document source format as are the full text sections and their headings.

## 3 Results

GePI is realized as a web application with a graphical user interface. The interface has two main levels: The **query level** and the **result level**. The opening page shows the options for querying specific gene/protein sets. In essence, two text fields enable the unrestricted specification of two lists of genes or proteins - termed *A*-list and *B*-list from here on. The input should either be NCBI Gene IDs, NCBI Gene symbols or FamPlex identifiers. The input to those two lists determine the topology of the search. Specifying only *A* items will result in an *open search*. This mode retrieves gene and protein interactions between the entries of *A* and any other interaction partners they have in the interaction database without restrictions. If a second list of gene identifiers is entered into the *B*-list, a *closed search* is performed. This mode is similar to the first but restricts the retrieved interaction items to those that have an element of *A* as one and an element of *B* as the other argument in any retrieved interaction description. It is of note that interactions between a single set of genes/proteins can be retrieved by performing a *closed search* with the identical lists queried as entries in *A* and *B*.

A number of NCBI Taxonomy IDs may be specified in an extra input field to restrict the search to genes or proteins that have been associated with one of the input organisms by GNormPlus or GeNo. This influences query processing as explained further below.

The input form at the **query level** offers input fields for the specification of full-text queries on the sentence or paragraph level. This enables defining queries that aim at a specific context scope and filter retrieved interactions accordingly. The input fields accept standard ES boolean queries (e. g. “X *OR* Y”) and are applied to filter the document context of the interaction items. Depending on the input field, the filter is applied on the sentence or the paragraph level. The sentence-level context is in close proximity to the linguistic structures that constitute the interaction statement and can thus be used to specify filter terms directly related to the interaction itself. The paragraph level widens the textual window around the interaction statement and allows to retrieve events with related context information beyond the sentence-scope. Of note, both textual scopes can be used complementary rather than alternatively. Consequently, both filter queries can be specified and connected by a boolean *AND* or *OR* operator to retrieve the description of the desired interaction context.

Similarly, another full-text filter can be applied for the section headings an interaction is mentioned in (e.g. restrict the search to “Results” sections only). Introductory and Background sections mostly refer to prior knowledge while Results sections reveal newly acquired insights described in the respective publication. Of note, heading types follow only rough naming conventions such as introduction, method or result sections and may vary. The possibility to include boolean queries allows for complex search requests that are able to tackle common section headings.

There are two ways to specify a GePI request. The first one is to enter at least one gene identifier into the *A*-list. The *B*-list is disabled as long as the *A*-field is empty. The second possibility is the usage of one of the full-text fields. If both *A* and *B* are empty, the results of this full-text-only-search comprise all molecular events in the interaction database that appear in a sentence, paragraph or section that matches the respective query filters. Hence, *A* and *B* serve as filters to a full-text query given for sentences, paragraphs or section headings. This query type can be used to identify published molecular interactions associated with a specific term, such as a condition (e. g. “elderly”) or a disease (e. g. obesity).

Once the query has been specified and sent to GePI, the gene identifiers are processed before the actual search is issued. By default, GePI resolves the input genes to their orthology surrogate concept as described in subsection 2.3 to retrieve results in an an organism-independent manner. If NCBI Taxonomy IDs have been specified in the respective input field, GePI will refrain from the ortholog resolution. Input gene IDs and gene symbols will then only be resolved to genes associated with one of the given taxonomy IDs.

After the successful query processing, the **result level** of GePI is presented. The retrieved interaction information is displayed in the form of several panels each of that focusses on a different aspect of the data to deliver a quick overview of the search result. The information shown in the panels falls into two general categories: aggregated overviews and the table of primary data where each retrieved interaction is listed individually with its textual context. The goal of the aggregation panels is to give a quick overview of the publication state with respect to the input. We use the frequency of genes, proteins and their pairwise interactions as a measure of literature-based support for the existence and scientific establishment of molecular events. Consequently, the majority of GePI’s result displays indicate a measure of occurrence frequency. The pie chart panel shows how often particular genes or proteins take part in any interaction that was returned for the current query. Association information about pairs of genes or proteins is depicted using Sankey diagrams. They connect symbols of genes or proteins that have been found to be associated in the literature with an edge of varying thickness. The thickness indicates the portion of interaction findings between two symbols. Stronger connections correlate with a higher frequency of identified interactions. There are two panels showing Sankey diagrams. The first shows the most frequent interactions in the retrieval result. The second Sankey diagram displays common interactions partners. It focusses on gene symbols that are frequently described to interact with a common, third symbol and thus shows second degree interactions. These results allow for finding new interactions, which is particularly meaningful for *closed search* requests, where non-trivial interactions might be present in the literature.

Finally, the table panel offers a complete list of associations between genes or proteins that have been found through the current request. It discloses detailed information about the gene mention in the text, the NCBI Gene ID and symbol it was mapped to, which algorithm(s) found the interaction partners and the sentence in which the interaction occurs. This is the primary result data from which all charts are derived.

The primary data table can be downloaded. This creates an Excel workbook that includes the details of the query and the resulting interaction data. The workbook also lists the exact occurrence numbers of the gene or protein symbols that take part in the interactions and how often each symbol was found in interaction with another symbol. The latter statistics are the background data of the pie and Sankey charts. The Excel workbook resembles the core output of a GePI session. It documents the query as well as the query result, provides further statistics to build upon and allows for efficient downstream analysis.

## 3.1 Evaluation

The result quality of GePI is largely determined by the document processing components. Most importantly, the gene recognition, gene normalization and event extraction algorithms have to produce high-quality output to maximize the number of correct interaction results and minimize false positives. Table 1 shows the scores reached by GNormPlus and GeNo on the BC2 and BioCreative III (BC3) GN challenge test data sets. All scores have been determined by evaluations carried out for this study with the exception of the BC2 scores for GNormPlus, which was trained on the whole of the BC2 GN dataset. The download version of the software prohibits re-training the GR model so we could not create a train-only model. Moreover, since gene mentions from the test set were added to the GNormPlus dictionaries, a comparable evaluation of the BC2 test data was not possible. Hence, we report here the official evaluation performance reported in Wei *et al*. (2015).

**Table 1.** Evaluation scores for the GN task on the BC2 and BC3 test data sets by GNormPlus and GeNo.

|  | BC2 Test |  |  | BC3 Test |  |  |
| --- | --- | --- | --- | --- | --- | --- |
|  | R | P | F | R | P | F |
| GNormPlus | 86.4% | 87.1% | 86.7% | 46.0% | 50.3% | 48.1% |
| GeNo | 84.0% | 81.4% | 82.7% | 21.2%* | 37.3%* | 27.1%* |

\*GeNo evaluation results on BC3 are exceedingly low due to data incompatibilities (*cf.* text description).

The performance values for BC3 in Table 1 were obtained by first converting the original PMC XML documents of the BC3 gold test set into the BioC (Comeau *et al*., 2013) format used by GNormPlus with our own conversion algorithm. Next, we processed the documents in BioC format, which is used to incorporate GNormPlus into the GePI production system. Thus, the displayed numbers reflect the output quality to be expected in the final system. This also explains the difference to the official numbers where GNormPlus is reported to reach an F-Score of 50.1. There might be small deviations between our BioC conversion and the document format used for the official GNormPlus evaluation.

Overall, GNormPlus compares favourably to our updated GeNo version with a 4% difference in F-Score on the BC2 test set. The gap on the BC3 test data is even larger potentially due to a data incompatibility issue: In current versions of NCBI Gene, 568 of the 1, 644 unique gene IDs that were present in the BC3 test set have been withdrawn due to re-sequencing processes in the database. Thus, the entries for these IDs were missing from our updated GeNo resources and cannot be found. For reference we provide the scores reached on the BC3 training set which is a recall of 43.8%, precision of 47, 2% and an F-Score of 45.4%. We did not use any of the BC3 data in the GeNo update for training and parameter tuning so that this is unseen data to our system. While these numbers cannot be used for direct comparison, the score on the training data showed that GeNo is functioning on full-text data in a multi-species setup.

The second essential step is the extraction of molecular events from the literature. The BioSem software we integrated into GePI has been evaluated by the authors for the BioNLP Shared Task 2011 and 2013 challenges on event extraction. We confirmed the originally reported performance scores reported in Bui and Sloot (2012) with our own experiments (Table 2). We opted for using BioSem to exploit its exceptional precision scores. Given the hardness of the event extraction task we opted for a high-precision system to optimize the correctness of identified interactions and consequently to avoid large and potentially false-positive rich result sets produced by GePI.

**Table 2.** Scores reached by BioSem in the BioNLP Shared Tasks 2011 and 2013 on the respective test data.

|  | R | P | F |
| --- | --- | --- | --- |
| BioNLP ST11 abstracts | 41.89% | 69.72% | 52.34% |
| BioNLP ST11 full texts | 44.47% | 66.63% | 53.34% |
| BioNLP ST13 | 42.47% | 62.83% | 50.68% |

## 4 Discussion

Automated molecular event extraction from scientific literature is able to provide gene or protein interaction information as soon as the respective papers are added to the literature databases. In doing so, it circumvents the need to query specific curated databases, which by definition cannot capture all of the literature describing molecular event information. Towards this aim we developed NLP modules(Hahn *et al*., 2016) that allowed us to extract all molecular event information from the biomedical literature at any given time point. We successfully applied our NLP efforts to identify potential interaction partners of proteins found in phosphoproteomics experiments including the AMPK complex (Dalle Pezze *et al*., 2016). We subsequently updated our retrieval algorithm to include full-text filter capabilities and used these to identify interactions between members of the Akt family and their pyruvate dehydrogenase kinase isoform 2 (PDK2)s in the context of cellular stress (Heberle *et al*., 2019). We also used our NLP expertise to gain the current state of potential interaction partners on kinases of the Pi3K/Akt signaling network as given in the scientific literature based on quantitative phosphoproteomics as input for our NLP service (Reimann *et al*., 2020). Finally, we successfully combined results from our NLP event extraction efforts with large-scale gene expression analysis and multiple validation experiments to generate a molecular signature of the activation of aryl hydrocarbon receptor (AHR) (Sadik *et al*., 2020). These examples demonstrated the flexible applicability of our NLP modules to complement life-science data set analysis and hypothesis generation. However, flexible usage of our NLP service without the necessity of expert knowledge in the NLP domain was not available. Appreciating the growing interest in automated event extraction and to facilitate our service also to the non-expert, we significantly updated and automated the process of event extraction. Most importantly, we quantified repeatedly requested tasks, optimized resource and computation time efficiency of our algorithms and implemented an automated update algorithm that ensures access to the current state of the literature at all times. We combined all relevant NLP tasks into a fully automated pipeline and created the web application GePI.

Despite the value offered by GePI for fully updated and automated event extraction, there is also room for improvement. On the preprocessing level, GNormPlus provides convincing performance but could possibly be enhanced by the integration of upcoming state-of-the art GR techniques. An updated algorithm combined with an enhanced species assignment algorithm to improve results on the BioCreative VI (BC6) Bio-ID assignment task was recently presented. Also, the aged resources of the tool might be inhibitive for some users. While we incorporated an updated GeNo version to alleviate a model based on outdated resources, the output quality could not be shown to outperform other gene recognition systems. BioSem was used for its superior specificity performance but has been surpassed by deep learning-based approaches that might identify additional interaction events. However, enhancements in these areas come with a pronounced increase in computational costs since modern multilevel neural networks pose high requirements on computing hardware, which is not suited for a production system such as we envision for GePI. In addition, given the long-term perspective of GePI, we developed our NLP pipeline in a component-based manner that allows easy exchange of individual text analysis components in the future.

## 5 Conclusion

In this work we presented GePI, a software tool for the fully automated extraction of molecular events from the biomedical literature and the retrieval of such events from an automatically populated database. The database includes interactions mined from the whole of PubMed and PubMed Central with regular updates to keep up with newest developments in the literature. We presented powerful query options for the retrieval of interactions between genes and proteins including an open search based on a single list of input gene identifiers and the possibility of a closed search where the potential interaction partner are specified beforehand. Together with options for full-text filter queries, GePI is highly adaptable to specific information needs. The interaction results can be downloaded as an Excel workbook that contains each found interaction and statistics about the frequency of the interaction partners and the interactions themselves. For each result interaction, its textual source is provided for reference. The fundamental NLP elements of GePI’s features were successfully used in multiple research scenarios that demonstrated its utility for biomolecular research. The here presented open access NLP pipeline and its access via a web application opens our service to the broad scientific community to complement and drive scientific discovery in the life-sciences.

